# Linker histone H1 determines cell stiffness and differentiation

**DOI:** 10.1101/2020.01.21.914770

**Authors:** Marie Kijima, Hiroyuki Yamagishi, Riho Kawano, Tomoki Konishi, Takuya Okumura, Masanori Hayase, Ryushin Mizuta

## Abstract

The flexibility or stiffness, one mechanical property of cells, is a promising label-free biomarker for underlying cytoskeletal or nuclear changes associated with various disease processes and changes in cell state. However, the molecular changes responsible for the whole-cell mechanical stiffness remain to be clarified. Recently, it was shown that the deterministic lateral displacement (DLD) microfluidic device, originally developed for size fractionation of some particles, might be applied to distinguish cells according to their stiffness. In this experiment, using the DLD device and various cell lines differentially expressing histone H1, a positively-charged protein localized in the linker region of chromatin, we found linear relationships between histone H1 quantity and cell stiffness. We also found that the histone H1 quantity affected cell size and even cell differentiation.

---

Histone H1, a positively-charged abundant nuclear protein, is located in the linker region of chromatin. Some in vitro studies using reconstituted chromatin showed that histone H1 tightly packs chromatin and results in a higher-order structure (9, 14, 28). Higher eukaryotes contain multiple duplicated *histone H1* genes. Both human and mouse have 11 *histone H1 variant* genes per haploid genome (7, 14). Null mutant cells have not been generated in human and mouse; however, in chicken, which has six *histone H1 variant* genes per haploid genome, one null mutant cell line (K11-5 or ΔΗ1) has been reported (8, 16). The ΔH1 cell was generated from chicken B-lymphoma DT40 by deleting all *histone H1 variant* genes.

Previously, using DT40 and ΔH1 cells, we showed that linker histone H1 was a chromatin condensation factor in both apoptotic and live cells, and without histone H1 chromatin became open (15, 16). Based on these findings, we proposed that histone H1 might determine the mechanical property of cells of flexibility (or stiffness). This property can be a useful marker of cell state and differentiation (3), although the responsible molecular mechanism has not been clarified. Recently, it was shown that the deterministic lateral displacement (DLD) microfluidic device, originally developed for size fractionation of some particles, might be applied to distinguish cells according to their stiffness (3, 10, 22).

In this experiment, we fabricated a similar DLD device and compared the cell stiffness among cell lines that showed different histone H1 expressions. We also examined the contribution of histone H1 to cell size and cell differentiation.

## Materials and Methods

### Plasmid construction and transfection

To construct an expression vector that carries a gene encoding a histone H1 protein, full-length coding sequences of chicken histone *H1R* were amplified from the pH1R-EGFP plasmid (8) using primers containing XhoI sites. The primer sequences used follow: cH1R-F2, 5′-CATG<u>CTCGAG</u>TGTTCGCAGCGCCGCCATGGCTG-3′; and cH1R-R2, 5′-GATC<u>CTCGAG</u>TTACTTCTTCTTGGCCGCTGCCTTCTTCG-3′. The amplified fragment was digested with XhoI and subcloned into the pPyCAGIRESzeocinpA expression vector, which consists of the CAG promoter, an internal ribosome entry site, the zeocin resistance gene, and a globin poly(A) site (19). The histone H1R expression vector was electroporated into a human histiocytic leukemia cell line U937 and zeocin-resistant clones UHZ-1 and UHZ-3 were established.

### Cell culture

Chicken B cell line DT40, linker histone H1-null mutant cell line ΔH1 (8, 16) were cultured in RPMI-1640 medium (Nissui Pharmaceutical Co., Tokyo, Japan) supplemented with 50 µM 2-mercaptetanol, 2 mM penicillin/streptomycin, 10% fetal calf serum (Thermo Fisher Scientific, Waltham, MA, USA), and 1% chicken serum (Nippon Bio-Test Laboratories Inc., Saitama, Japan) at 40°C under 5% CO2 in a humidified incubator. Cell lines U937, UHZ-1, and UHZ-3 were cultured in RPMI-1640 medium supplemented with 50 µM 2-mercaptetanol, 2 mM penicillin/streptomycin, and 10% fetal calf serum at 37°C under 5% CO_2_ in a humidified incubator.

### Western blot analysis

Cells (1.0 × 10^6^) were harvested, suspended in 50 µL of phosphate buffered saline (PBS) and lysed with 50 µL of sample buffer [4% sodium dodecyl sulfate (SDS), 100 mM Tris-HCl (pH 6.8), 2 mg/mL bromophenol blue, 20% glycerol, and 200 mM dithiothreitol]. After 5 min of sonication, the samples were separated on an SDS-15% polyacrylamide gel and then electrophoretically transferred onto the nitrocellulose membrane Amersham Protran 0.45 µm NC (GE Healthcare Lifesciences, Chicago, IL, USA). Membrane-immobilized proteins were probed with mouse anti-linker histone H1 monoclonal antibody clone AE-4 (Upstate Biotechnology, Lake Placid, NY, USA) or goat anti-actin antibody clone I-19 (Santa Cruz Biotechnology, Dallas, TX, USA) and visualized with horseradish peroxidase-conjugated rabbit anti-mouse IgG (Thermo Fisher Scientific) or horseradish peroxidase-conjugated donkey anti-goat IgG (Santa Cruz Biotechnology), respectively. The AE-4 antibody cross-reacts with core histone H3 (17). The signal of histone H3 or actin was used as the loading control. Images were collected with LAS-3000 Imager and analyzed with Image Gauge software (GE Healthcare Lifesciences).

### Microfluidic DLD device

Detail of the fabrication of DLD device was shown in Supplemental Materials and Methods. The DLD post array dimensions were determined by preliminary experiments (Fig. 1a). Cells were suspended in the buffer solution (RPMI-1640 supplemented with 12.5 mM HEPES and 0.5% chicken serum) and the concentration adjusted to 3.0 × 10^5^ cells/mL. Solutions were injected into the micro-channel by syringe pumps. A dispersed cell solution was poured into the sample inlet, and buffer solution was poured into the buffer inlet. The ratio of cell to buffer was 1:25. The sample solution was surrounded by buffer solution, and a thin stream-line which entered the micro-posts array zone was formed. In this study, four configurations of different fluid velocities were tested: Slow, Normal, Fast 1, and Fast 2 of 50, 100, 300, and 600 µL/min, respectively (Fig. 1b). Separation behavior was recorded with a FASTCAM SA3 high-speed camera (Photron, Tokyo, Japan), and the cell number and the length displaced from the position of the inlet nozzle (lateral displacement length, LDL) were determined. The average LDL was calculated using the following equation in each flow condition.

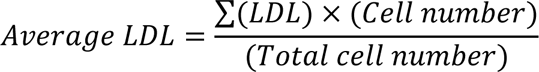

**Fig. 1.**
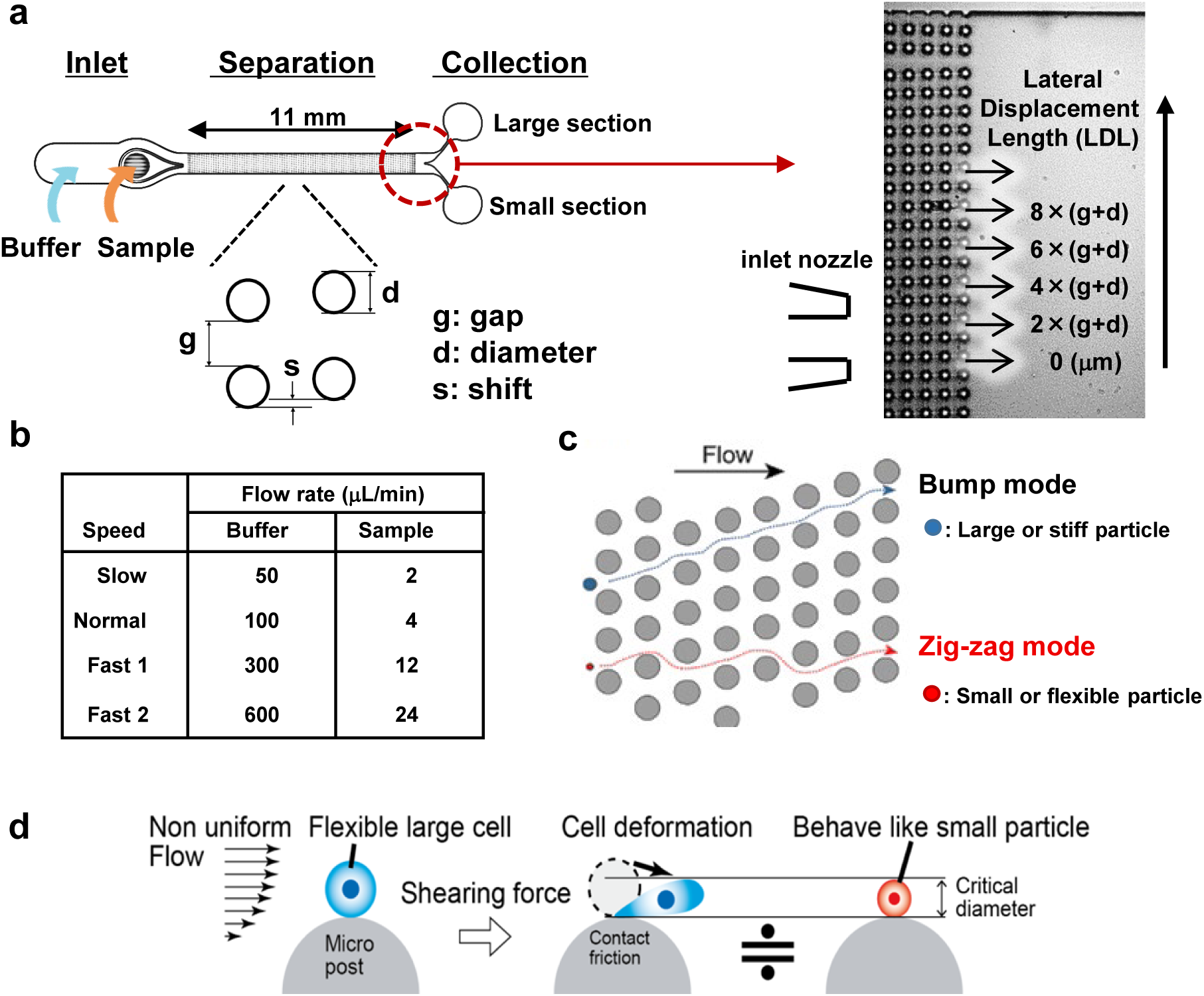
Microfluidic device and deterministic lateral displacement (DLD). (a) The DLD device used in this experiment had three parts: inlet, separation, and collection. In the separation part, small columns were distributed regularly for separating cells and the dimensions, such as gap (g), diameter (d), and shift (s), were determined by preliminary experiments. Separation behavior was recorded with a high-speed camera, and cell number and positions of cells at the exit of separation zone, indicated with red dotted circle, were determined. The magnified image of the exit along with the position of inlet nozzle was shown in the right. Lateral displacement length (LDL) of a cell was determined with the number of column portion, gap (g), and diameter (d), and the average LDL was calculated from cell number and LDL. (**b**) Four configurations of different fluid velocities are shown. Using syringe pumps, dispersed sample suspension and buffer are injected into the sample and buffer inlets, respectively. The sample solution is surrounded by buffer solution and a thin stream-line is formed. (**c**) Separation behavior of particles. In the micro-post array zone, laminar flow is expected and no turbulence happens. Large or stiff particles are displaced laterally after collisions with columns, and flow into the large section (bump mode). Small or flexible particles are deformed after collisions with columns, and flow into the small section (zig-zag mode). (**d**) Principle of DLD. Flexible cells are deformed in the flowing condition and behave as small particles.

Statistical analysis was conducted using data from three independent experiments (Supplemental Fig. S1 and S2).

### FACS analysis

Cells (4.0 × 10^6^) were harvested and rinsed with PBS. Then, cells were suspended in FACS buffer (0.5% BSA, 2 mM EDTA, and 0.005% sodium azide in PBS) and analyzed by BD FACSCanto II (Becton, Dickinson and Company, Franklin Lakes, NJ, USA). Forward scatter (FSC) and side scatter (SSC) were the parameters for selecting viable cells.

### Cell differentiation

Cells (2.0 × 10^6^ or 4.0 × 10^6^) were stimulated with phorbol 12-myristate 13-acetate (PMA, 2 mg/mL). Cells were harvested at indicated times after stimulation. Some cells were stained with biotin-labeled anti-CD11b antibody (BioLegend, San Diego, CA, USA) and streptavidin–fluorescein isothiocyanate conjugate (Thermo Fisher Scientific), and analyzed by BD FACSCanto II (Becton, Dickinson and Company).

### 2.7. Quantitative PCR (q-PCR) analysis

The *p21* expression was confirmed by q-PCR analysis. The expression of glyceraldehyde-3-phosphate dehydrogenase (GAPDH) was used as the control. Total RNA was extracted from cells using TRIzol (Sigma Aldrich, St. Louis, MO, USA) according to the manufacturer’s instructions. The RNA (10 μg) was reverse-transcribed into cDNA using Superscript II with oligo (dT) primers (Thermo Fisher Scientific). The cDNA was amplified in a thermal cycler 7500 Fast (Applied Biosystems, Foster City, CA, USA) with THUNDERBIRD SYBR qPCR Mix (Toyobo, Osaka, Japan) and primers. The primers used follow: GAPDH-F, 5′-GACCTGCCGTCTAGAAAAAC-3′;

GAPDH-R, 5′-TTGAAGYCAGAGGAGACCAC-3′; p21-F,

5′-CTGGAGACTCTCAGGGTCGAA-3′; and p21-R,

5′-GGATTAGGGCTTCCTCTTGGA-3′.

## Results

### Microfluidic device to separate cells by cell stiffness

We fabricated a DLD device and compared cell stiffness between histone H1-sufficient and -deficient cell lines. The device was composed of three parts: inlet, separation, and collection (Fig. 1a). In the separation part, a periodic array of micro-posts, shifted slightly along the fluid stream, was fabricated. The dimensions of the post array – gap (g), diameter (d), and shift (s) – were determined in each experiment. Four configurations of flow rates of buffer and sample are shown in Fig. 1b. In the microfluidic channel, laminar flow without turbulence was expected (10, 22). Small particles follow the fluid stream-line and travel in a “zig-zag mode”, while large particles bump on the posts, travel across the fluid stream-line, and flow at a slight angle along the post array in a “bump mode” (Fig. 1c). In DLD theory, the critical separation diameter can be estimated assuming a rigid sphere (12), but shear stress is applied to particles by non-uniform fluid velocity or contact friction on the micro-posts. Flexible particles are deformed in the flowing condition and behave as smaller particles (Fig. 1d). Therefore, it is assumed that cell rigidity can be determined from the separation behavior. We took video images of cells flowing out of the separation zone and counted cells exited from each lateral displaced portion (Fig. 1a, right). The average lateral displacement length (LDL) calculated from the cell number and the length displaced from the position of the inlet nozzle is the parameter of deformation or stiffness of cells in each flow speed (Supplemental Fig. S1 and S2).

### Loss of histone H1 reduced cell stiffness in DT40 cells

We compared DT40 and ΔH1 cells, where the latter cells had all *histone H1* genes deleted from the DT40 genome. First, we confirmed the loss of histone H1 in ΔH1 by western blot analysis (Fig. 2a), and also determined cell size by flow cytometry (Fig. 2b). As expected, ΔH1 cells were slightly larger than DT40 cells, because Hashimoto et al. showed that ΔH1 had more expanded nuclei than DT40 (8), suggesting that the loss of histone H1 increased cell size. Then, we ran DT40 or ΔH1 cells in the microfluidic device (Fig. 2c) under four flow speed conditions: Slow, Normal, Fast 1, and Fast 2 (Fig. 1b). We counted cells exited from different displaced sites and determined the average LDL (Supplemental Fig. S1). If DT40 and ΔH1 had the same cell stiffness, ΔH1 would be displaced more than DT40, because ΔH1 is larger than DT40 (Fig. 2b). However, our results revealed greater displacement length in DT40 than ΔH1 cells at each flow speed (Fig. 2d), suggesting that ΔH1 was more flexible than DT40. This result indicated that the loss of histone H1 decreased cell stiffness.

**Fig. 2.**
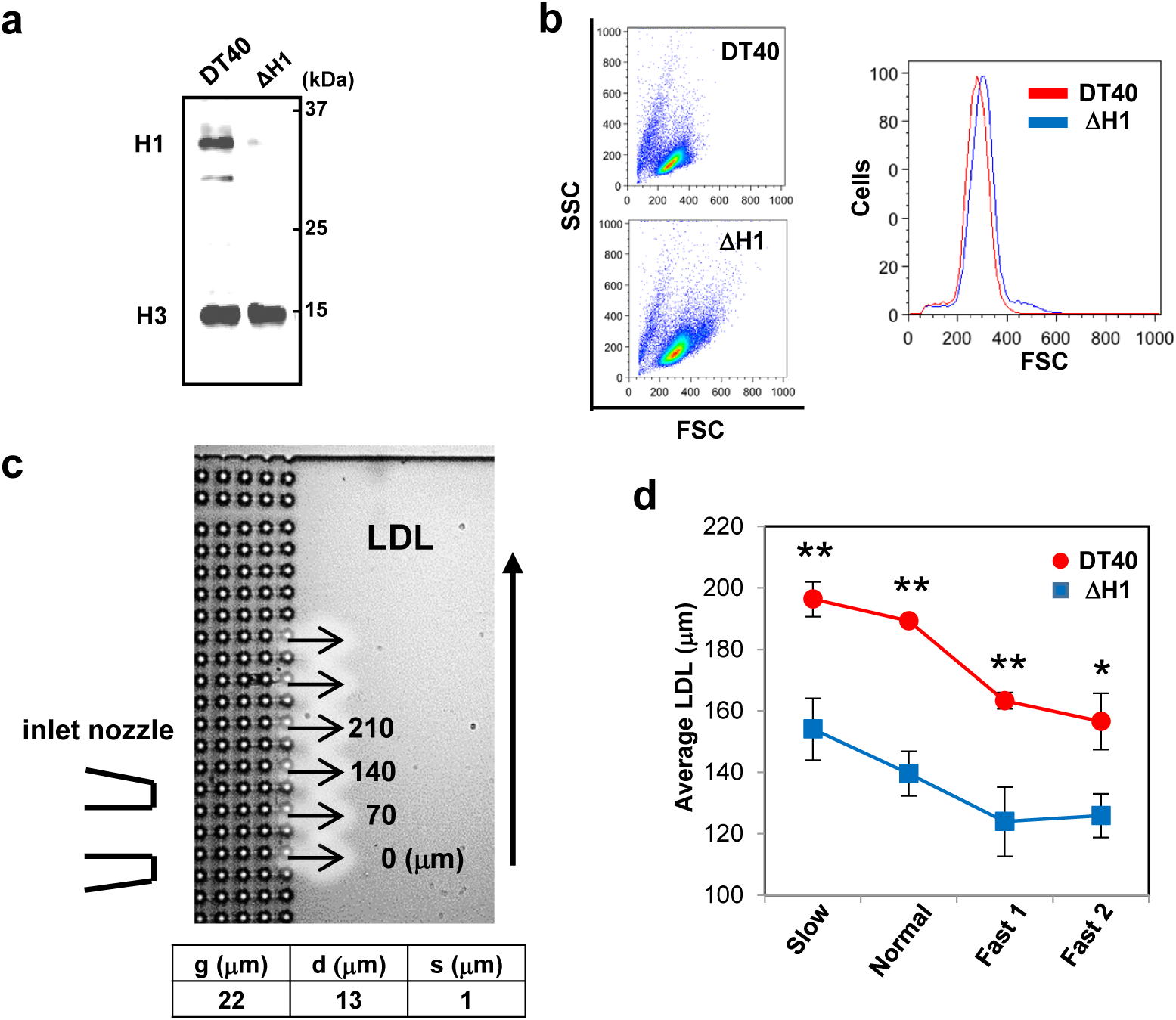
Loss of histone H1 reduced cell stiffness in DT40 cells. (a) Western blot analysis of histone proteins. Histone H1 (H1) was deleted in ΔH1 cells. Histone H3 (H3) was a loading control. (**b**) Flow cytometric analysis showed that ΔH1 cells were larger than DT40 cells. The forward scatter (FSC) and side scatter (SSC) (left), and FSC histograms of viable DT40 and ΔΗ1 cells are shown (right). (**c**) Dimensions of the microfluidic device and portions of lateral displacement. (**d**) The ΔH1 cells were more flexible than DT40 cells. The DT40 or ΔH1 cells were separated in the microfluidic device under four flow speed conditions: Slow, Normal, Fast 1, and Fast 2 (Fig. 1b). Separated cells were counted and the average LDL calculated (Supplemental Fig. S1). Data are shown as mean ± SD of three independent experiments. *, *P* < 0.05; **, *P* < 0.01 (two-tailed unpaired *t*-test).

### Increased histone H1 augmented cell stiffness and chromatin condensation in U937 cells

To confirm the above results, we initially intended to express one of the chicken *histone H1* genes in ΔH1. However, in the process of generating ΔH1, seven antibiotic genes were already introduced (8, 26) and we could not establish another stable transformant. Consequently, we focused on the human histiocytic leukemia cell line U937, because it was reported to have a lower histone H1 quantity (11, 15). The homology of histone H1 proteins is high and their functions are conserved beyond species (23). Thus, we introduced a chicken histone H1 expression vector and generated two stable clones: UHZ-1 and UHZ-3. The histone H1 expression was determined by western blot analysis (Fig. 3a). Exogenous chicken histone H1 that was slightly smaller than endogenous human H1 was detected in both UHZ-1 and UHZ-3. Flow cytometry showed that UHZ-1 and UHZ-3 had similar cell size and were smaller than U937 (Fig. 3b), suggesting that UHZ-1 and UHZ-3 became more compact with histone H1 overexpression. Then, we flowed UHZ-1 and UHZ-3 through the microfluidic device (Fig. 3c) and found that both lines behaved similarly and tended to be more laterally displaced than U937 (Fig. 3d and Supplemental Fig. S2). Notably, UHZ-1 and UHZ-3, despite being smaller than U937 cells, had significantly longer displacement length than U937 at the Fast 1 flow speed, suggesting that they were stiffer than U937. Thus, histone H1 quantity might be linearly correlated with cell compactness and stiffness. Previously, using DT40 and ΔH1, we showed that histone H1 quantity determined the efficiency of chromatin condensation in both apoptotic and live cells (16), and then we aimed to confirm this finding using U937 and UHZ-1 cells. First, we induced apoptosis by ultraviolet irradiation and found more chromatin condensation in UHZ-1 than U937 (Supplemental Fig. S3a). Next, we compared the DNase accessibility to isolated nuclei. We incubated isolated nuclei from U937 or UHZ-1 with one of three endonucleases: micrococcal nuclease (MNase), DNase I, and DNase γ (also named DNase IL3). We found reduced chromatin degradation of UHZ-1 nuclei compared to U937, and longer DNA fragments were detected (Supplemental Fig. S3b). These results suggested that cells containing more histone H1 had more condensed chromatin, similar to the previous results using DT40 and ΔΗ1 cells.

**Fig. 3.**
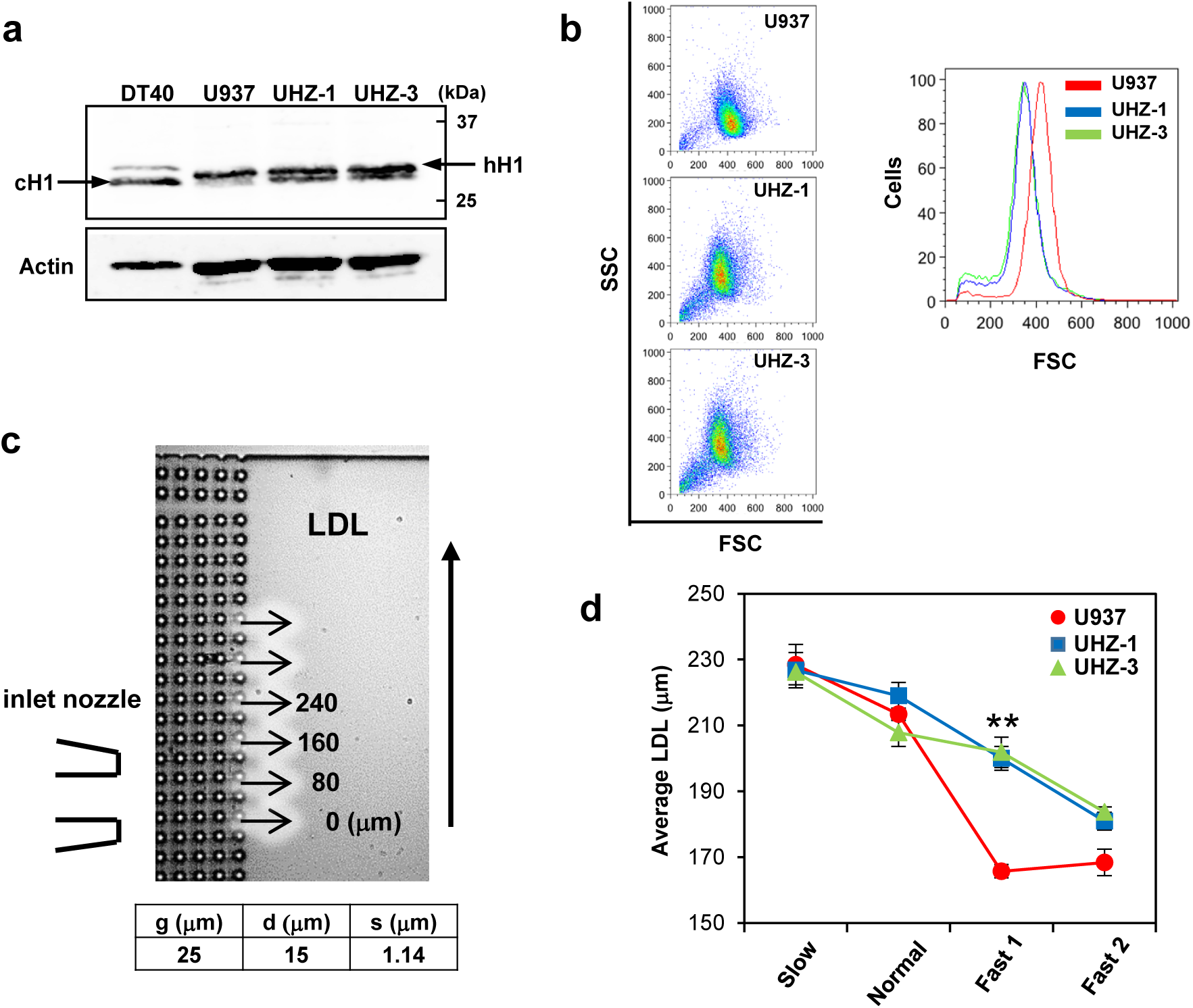
Increased histone H1 augments cell stiffness in U937 cells. (a) Western blot analysis of histone proteins in DT40, U937, UHZ-1, and UHZ-3 cells. Exogenous chicken histone H1 (cH1) and endogenous human histone H1 (hH1) are shown. Actin was a loading control. (**b**) Flow cytometric analysis showed that UHZ-1 and UHZ-3 had similar cell sizes and were smaller than parental U937. The FSC and SSC analysis (left) and FSC histograms are shown (right). (**c**) Dimensions of the microfluidic device and portions of lateral displacement. (**d**) UHZ-1 and UHZ-3 were stiffer than U937. Cells were separated in the microfluidic device under four flow speed conditions: Slow, Normal, Fast 1, and Fast 2 (Fig. 1b). Separated cells were counted and the average LDL calculated (Supplemental Fig. S2). Data are shown as mean ± SD of three independent experiments (U937 and UHZ-1) and two independent experiments (UHZ-3). **, *P* < 0.01 (U937 vs UHZ-1, two-tailed unpaired *t*-test).

### Histone H1 quantity affects U937 differentiation

Cell line U937 can be induced to differentiate to a macrophage-like cell (5, 18). Thus, we wanted to know how overexpression of histone H1 could affect the differentiation of U937. We induced differentiation of U937, UHZ-1, and UHZ-3 to macrophage-like cells using PMA, and cell growth was significantly inhibited in UHZ-1 and UHZ-3 (Fig. 4a). To confirm this finding, we also examined the expression of cell cycle regulator p21, a cyclin-dependent kinase inhibitor capable of inhibiting all cyclin/cyclin-dependent kinase complexes (6). As expected, expression of *p21* was significantly higher in UHZ-1 and UHZ-3 than U937 (Fig. 4b). Differentiation of U937 was determined by flow cytometry after staining a cell surface marker of macrophage CD11b (Fig. 4c) (5). There was higher expression of CD11b and more differentiation to macrophage-like cells in UHZ-1 and UHZ-3 than U937 (Fig. 4d). Thus, histone H1 quantity affected U937 differentiation.

**Fig. 4.**
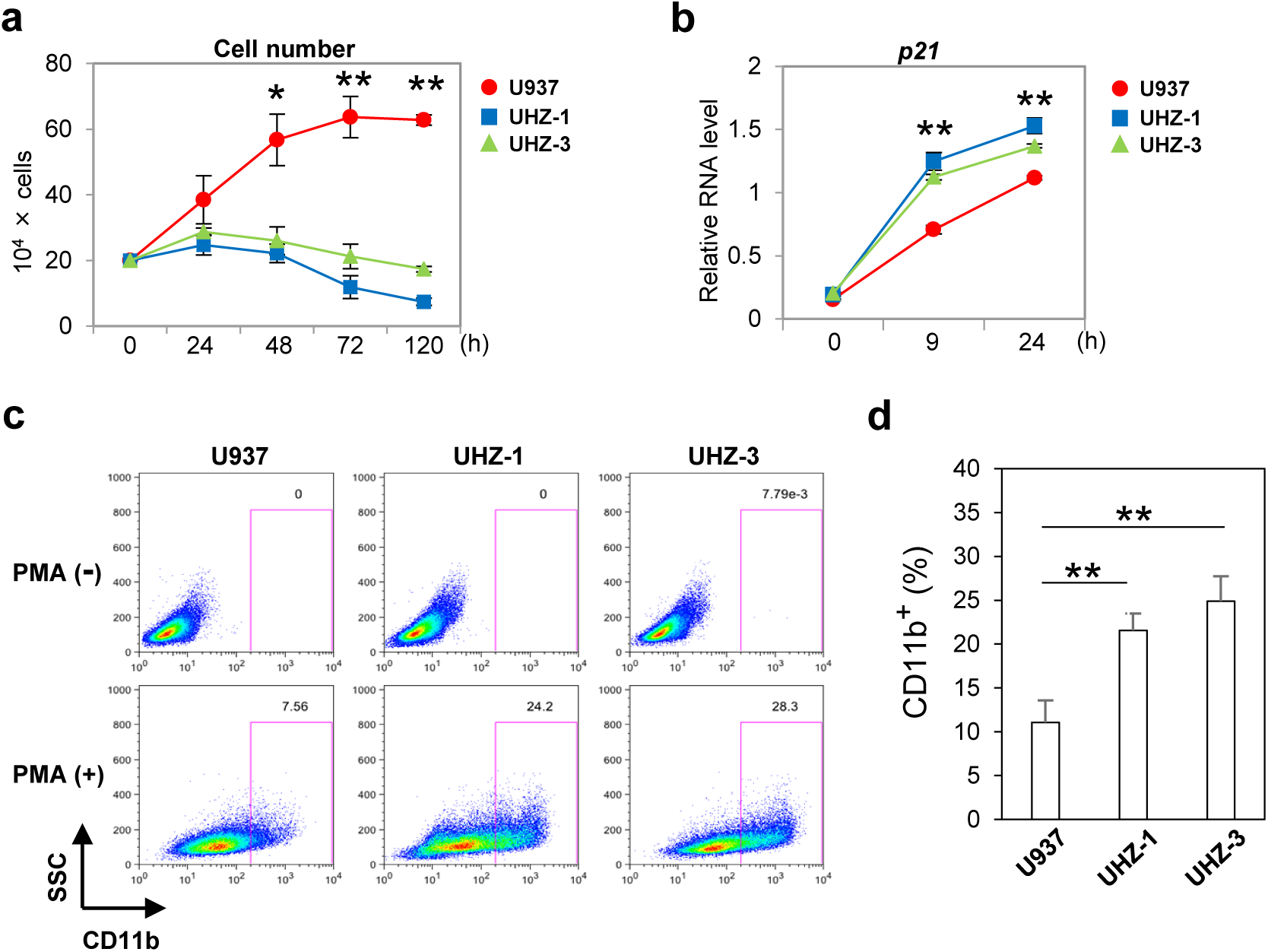
Histone H1 quantity affects U937 differentiation. (a) Cells (2.0 × 10^6^) were stimulated with PMA and cell growth was determined at 0–120 h. Data are shown as mean ± SD of three independent experiments. *, *P* < 0.05, **, *P* < 0.01 (U937 vs UHZ-1 or UHZ-3, two-tailed unpaired *t*-test). (**b**) Relative mRNA levels of *p21* at each incubation time, 0–24 h. Data are shown as mean ± SD of three independent experiments. **, *P* < 0.01 (U937 vs UHZ-1 or UHZ-3, two-tailed unpaired *t*-test). (**c**) Flow cytometric analysis of cell differentiation to macrophage-like cells after 72 h of PMA stimulation. The CD11b expression was determined before and after PMA stimulation: PMA (-) and PMA (+), respectively. Representative images were shown. (**d**) Histograms show the ratio of CD11b positive cells after PMA stimulation. Data are shown as mean ± SD of three independent experiments. **, *P* < 0.01 (U937 vs UHZ-1 or UHZ-3, two-tailed unpaired *t*-test).

## Discussion

Cell stiffness, one mechanical property of cells, is a promising label-free biomarker for underlying cytoskeletal or nuclear changes associated with various disease processes and changes in cell state (3). For example, increased cell flexibility has been identified in pluripotent stem cells (20). Mouse and human embryonic stem (ES) cells and their nuclei were found to be more flexible than those of their differentiated progeny, suggesting flexibility may be a viable biomarker for pluripotency (1, 20). Furthermore, cell flexibility of metastatic and malignant cancer cells increased compared to benign cells of the same origin (24, 25). Interestingly, it was also reported that chemotherapy changed tumor cell flexibility (24). Thus, a simple technique for measuring cell stiffness may have clinical implications in screening cells for resistance and sensitivity to chemotherapeutic drugs. Additionally, activated lymphocytes become softer compared to naïve lymphocytes (3). Contrary to this, neutrophils and monocytes become stiffer upon stimulation with chemokines associated with infection (4, 21). Thus, flexibility or stiffness is a fundamental physical property that determines the state and character of cells, e.g., ES, cancer, and immune cells.

The molecular changes responsible for whole-cell mechanical stiffness remains to be clarified, although nuclear architecture, epigenetic state, and tighter chromatin packing are all suggested to affect cell stiffness (3, 20). In the process of differentiation, cells gain regions of condensed heterochromatin, which decreases cell flexibility and induces cell stiffness. However, global chromatin decondensation is required to maintain pluripotency and citrullination of linker histone H1 and its displacement from chromatin by peptidyl arginine deiminase type IV (PADI4) is essential for this process (2). Histone H1 is a heterochromatin protein and maintains chromatin condensation (13, 16, 27). Thus, it is reasonable to expect that histone H1 regulates cell stiffness. In this report, we showed that histone H1-deficient ΔH1 cells were larger and more flexible than the parental cell line DT40, confirming that histone H1 contributed to cell size and stiffness. Using U937 and its stable transformants overexpressing histone H1, we also showed that the change in histone H1 quantity affected cell size and flexibility, and even cell differentiation.

Higher eukaryotes contain multiple duplicated *histone H1* genes. Both human and mouse have 11 *histone H1 variant* genes and chicken has six per haploid genome (7, 14). Each variant protein has high homology, e.g., percentage identity of human H1.4 to human H1.1 or to chicken H1R is comparable – with 72.50% and 69.58%, respectively – suggesting that its function is conserved beyond species (23).

In this report, we used a DLD device to determine cell stiffness. Several devices and techniques have been developed to measure mechanical properties of cells, e.g., atomic force microscopy, micropipette aspiration, cell transit analyzer, and optical stretching (3). However, these approaches have disadvantages in that they are low throughput (<1 cell/min), measure only a part of cell, and sometimes require skilled manual operation. In contrast, the DLD device has advantages of high throughput and determining whole-cell mechanical property. Thus, the DLD device used in this experiment was suitable for comparing cell stiffness of DT40 vs ΔH1, or U937 vs UHZ-1 and UHZ-3. However, there some problems to overcome in the current DLD device: (1) it requires preliminary experiments to determine post array dimensions, (2) mixing a sample solution is required, and (3) occlusion sometimes occurs. Despite these problems, we expect that DLD devices will be useful and next generation DLD devices are expected. As mentioned above, several systems or devices to detect cell flexibility or stiffness have been developed (3). One problem of these systems is the lack of good standard cell lines for evaluation. Given that ΔΗ1 cells were generated from DT40 cells using an ordinary gene knock-out approach (8), we expect that DT40 and ΔΗ1 cells are useful as potential standard cell lines.

In this report, we showed that linker histone H1, a chromatin condensation factor, determined cell size and stiffness, and also cell differentiation. The physiological significance of histone H1 will be clarified further in future. We also hope that cell stiffness will be accepted as a standard biomarker of cell state and applied in clinical fields.

## Acknowledgements

This work was supported by Grants-in Aid for Scientific Research (C) 17K08891 from the Ministry of Education, Culture, Sports, Science and Technology (MEXT), Japan (to RM).

