## Supplementary material for "Linker histone H1 determines cell stiffness and differentiation": Supplymental materials

### Supplementary Data

### Materials and Methods

*Microfluidic DLD device*

The microfluidic device was fabricated using silicon substrate by conventional photolithography and the Deep Reactive Ion Etching (DeepRIE) process. Photoresist was spin-coated on the 550-μm thick silicon substrate with 500-nm thick thermal silicon oxide and DLD micro-post array pattern was transferred on the thermal oxide layer by wet etching with a hydrofluoric acid solution. Using the thermal oxide layer as a mask, silicon was etched about 50 μm by SPP MUC-21 (Sumitomo Precision Products, Tokyo, Japan) and the micro-post array was fabricated. After the DeepRIE process, the silicon substrate was dipped in a hydrofluoric acid solution to remove all silicon oxide. Prior to experiments, the silicon substrate was carefully cleaned and a thermal oxide layer was formed again to obtain hydrophilic surface. A glass cover plate with inlet and outlet holes was put on the silicon substrate and the microfluidic device was realized (Fig. 1a).

*Apoptosis induction*

To induce apoptosis, cells ($5.0 \times 10^5$ cells/mL) were irradiated with ultraviolet (UV) (40 mJ/cm$^2$) using GS Gene Linker UV chamber (Bio-Rad Laboratories, Hercules, CA, USA) and incubated at 37 °C. Condensed chromatin was stained with Hoechst 33342 (1 μg/mL) (Dojindo, Kumamoto, Japan) at 6 h after UV-irradiation and examined with an Olympus IX71 fluorescence microscope (Olympus, Tokyo, Japan).

*DNA fragmentation*

DNA fragmentation assay using isolated nuclei was described previously (1, 2). The reaction mixture for the DNA fragmentation assay in a final volume of 60 μL contained $2.0 \times 10^5$ nuclei and DNase — MNase (0.2 U), DNase I (0.02 U) or DNase γ (0.5 ng). The reactions were performed at 37 °C for 2 h. After incubation, the reaction samples were lysed by the addition of 300 μL of 8 M guanidine and DNA was isolated from the lysate with the Wizard DNA purification resin (Promega, Madison, WI, USA) as described. DNA was analyzed by electrophoresis on a 1.5% agarose gel in the presence of ethidium bromide (0.5 μg/mL).

a

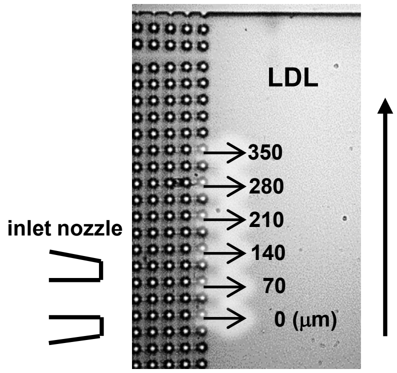


b

| Cell | DT40 | | | | | | | | | | | |
|---|---|---|---|---|---|---|---|---|---|---|---|---|
| Experiments | 716-1 | | | | 1220 | | | | 1222 | | | |
| LDL (μm) \ Speed | Slow | Normal | Fast 1 | Fast 2 | Slow | Normal | Fast 1 | Fast 2 | Slow | Normal | Fast 1 | Fast 2 |
| 350 | 0 | 0 | 0 | 0 | 1 | 0 | 0 | 0 | 0 | 0 | 0 | 0 |
| 315 | 0 | 0 | 0 | 0 | 7 | 3 | 0 | 0 | 11 | 5 | 3 | 3 |
| 280 | 6 | 8 | 2 | 4 | 8 | 5 | 3 | 2 | 25 | 14 | 11 | 9 |
| 245 | 19 | 18 | 20 | 20 | 22 | 37 | 22 | 11 | 15 | 14 | 2 | 5 |
| 210 | 22 | 22 | 35 | 27 | 53 | 42 | 29 | 25 | 39 | 37 | 22 | 24 |
| 175 | 13 | 13 | 24 | 25 | 32 | 24 | 36 | 41 | 42 | 70 | 51 | 48 |
| 140 | 15 | 17 | 41 | 35 | 15 | 26 | 33 | 43 | 15 | 16 | 39 | 32 |
| 105 | 6 | 12 | 32 | 20 | 2 | 8 | 17 | 19 | 4 | 2 | 12 | 16 |
| 70 | 3 | 2 | 5 | 7 | 6 | 2 | 22 | 17 | 5 | 3 | 9 | 14 |
| 35 | 1 | 1 | 0 | 3 | 8 | 9 | 4 | 9 | 3 | 5 | 8 | 2 |
| 0 | 0 | 0 | 0 | 0 | 0 | 0 | 1 | 4 | 1 | 1 | 0 | 0 |
| Total cell № | 85 | 93 | 159 | 141 | 154 | 156 | 167 | 171 | 160 | 167 | 157 | 153 |
| Average LDL (μm) | 190.6471 | 186.6667 | 166.4151 | 167.8014 | 194.3182 | 188.2372 | 159.9102 | 147.3684 | 204.0938 | 190.2994 | 163.6306 | 165.6209 |

| Cell | ΔH1 | | | | | | | | | | | |
|---|---|---|---|---|---|---|---|---|---|---|---|---|
| Experiments | 716-2 | | | | 1208 | | | | 1119 | | | |
| LDL (μm) \ Speed | Slow | Normal | Fast 1 | Fast 2 | Slow | Normal | Fast 1 | Fast 2 | Slow | Normal | Fast 1 | Fast 2 |
| 350 | 0 | 0 | 0 | 0 | 0 | 0 | 0 | 0 | 2 | 1 | 1 | 0 |
| 315 | 0 | 0 | 0 | 0 | 0 | 0 | 0 | 0 | 0 | 1 | 2 | 4 |
| 280 | 0 | 0 | 0 | 0 | 0 | 1 | 1 | 1 | 15 | 1 | 6 | 11 |
| 245 | 2 | 0 | 0 | 0 | 3 | 5 | 2 | 7 | 29 | 6 | 14 | 16 |
| 210 | 8 | 13 | 2 | 7 | 38 | 15 | 13 | 31 | 26 | 14 | 13 | 33 |
| 175 | 14 | 21 | 28 | 9 | 56 | 26 | 28 | 29 | 73 | 42 | 50 | 85 |
| 140 | 5 | 10 | 23 | 25 | 65 | 47 | 58 | 68 | 52 | 41 | 33 | 62 |
| 105 | 3 | 10 | 25 | 27 | 38 | 18 | 28 | 60 | 40 | 34 | 64 | 83 |
| 70 | 2 | 7 | 14 | 10 | 37 | 34 | 32 | 42 | 20 | 8 | 32 | 31 |
| 35 | 0 | 6 | 11 | 7 | 10 | 12 | 26 | 41 | 12 | 8 | 10 | 42 |
| 0 | 2 | 0 | 0 | 0 | 0 | 0 | 15 | 2 | 0 | 0 | 2 | 9 |
| Total cell № | 36 | 67 | 103 | 85 | 247 | 158 | 203 | 281 | 269 | 156 | 227 | 376 |
| Average LDL (μm) | 160.4167 | 142.6119 | 121.6505 | 121.4706 | 139.8583 | 129.5886 | 111.3793 | 120.1957 | 161.8587 | 146.5064 | 138.7665 | 135.9043 |

**Supplemental Figure S1**

**Number and displaced length of cells separated by a microfluidic device. (a)** Magnified image of the exit of separation area, the position of inlet nozzle, and portions of lateral displacement. **(b)** Cell number and lateral displacement length (LDL) at each flow speed were shown. Separated cells were counted and the average LDL calculated (Materials and Methods). Three independent experiments were conducted in both DT40 and ΔH1.

a

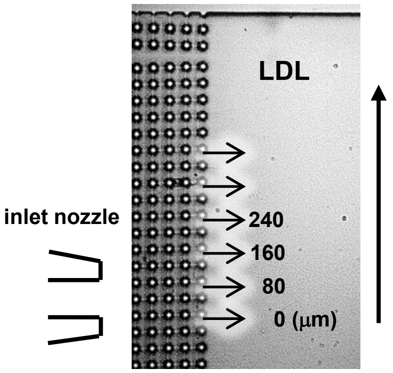


b

| Cell | U937 | | | | | | | | | | | |
|---|---|---|---|---|---|---|---|---|---|---|---|---|
| Experiments | 1216-1 | | | | 109-1 | | | | 112-1 | | | |
| LDL (μm) \ Speed | Slow | Normal | Fast 1 | Fast 2 | Slow | Normal | Fast 1 | Fast 2 | Slow | Normal | Fast 1 | Fast 2 |
| 360 | 0 | 0 | 0 | 0 | 0 | 0 | 0 | 0 | 6 | 4 | 0 | 0 |
| 320 | 0 | 1 | 0 | 0 | 1 | 0 | 0 | 0 | 7 | 3 | 0 | 0 |
| 280 | 7 | 1 | 0 | 0 | 11 | 11 | 8 | 0 | 3 | 7 | 2 | 0 |
| 240 | 69 | 61 | 17 | 35 | 25 | 31 | 55 | 50 | 3 | 9 | 41 | 173 |
| 200 | 31 | 25 | 39 | 17 | 10 | 23 | 25 | 183 | 26 | 22 | 21 | 84 |
| 160 | 12 | 13 | 40 | 26 | 3 | 11 | 80 | 159 | 6 | 9 | 46 | 201 |
| 120 | 2 | 5 | 30 | 40 | 1 | 8 | 44 | 115 | 2 | 8 | 21 | 185 |
| 80 | 2 | 0 | 5 | 4 | 1 | 1 | 28 | 47 | 0 | 0 | 14 | 17 |
| 40 | 0 | 1 | 2 | 1 | 0 | 0 | 13 | 9 | 0 | 0 | 11 | 10 |
| 0 | 0 | 0 | 0 | 0 | 0 | 0 | 0 | 2 | 0 | 0 | 1 | 0 |
| Total cell № | 123 | 107 | 133 | 123 | 52 | 85 | 253 | 565 | 53 | 62 | 157 | 670 |
| Average LDL (μm) | 219.8374 | 214.5794 | 168.1203 | 171.7073 | 232.3077 | 210.8235 | 163.1621 | 162.7611 | 233.2075 | 214.8387 | 165.8599 | 170.806 |

| Cell | UHZ-1 | | | | | | | | | | | |
|---|---|---|---|---|---|---|---|---|---|---|---|---|
| Experiments | 1216-2 | | | | 109-2 | | | | 112-2 | | | |
| LDL (μm) \ Speed | Slow | Normal | Fast 1 | Fast 2 | Slow | Normal | Fast 1 | Fast 2 | Slow | Normal | Fast 1 | Fast 2 |
| 360 | 0 | 0 | 1 | 0 | 1 | 4 | 0 | 0 | 2 | 2 | 1 | 8 |
| 320 | 0 | 5 | 2 | 0 | 8 | 8 | 0 | 1 | 19 | 13 | 8 | 8 |
| 280 | 8 | 42 | 37 | 9 | 2 | 8 | 1 | 0 | 2 | 10 | 6 | 24 |
| 240 | 14 | 19 | 16 | 29 | 7 | 26 | 35 | 21 | 27 | 49 | 16 | 46 |
| 200 | 1 | 20 | 20 | 9 | 18 | 31 | 6 | 44 | 27 | 24 | 17 | 96 |
| 160 | 4 | 21 | 40 | 38 | 5 | 16 | 16 | 35 | 18 | 23 | 24 | 104 |
| 120 | 1 | 5 | 12 | 9 | 0 | 5 | 5 | 6 | 6 | 13 | 8 | 15 |
| 80 | 0 | 5 | 5 | 9 | 3 | 4 | 5 | 7 | 2 | 3 | 3 | 42 |
| 40 | 0 | 0 | 1 | 1 | 0 | 1 | 0 | 4 | 0 | 0 | 2 | 14 |
| 0 | 0 | 0 | 0 | 0 | 0 | 0 | 0 | 0 | 0 | 0 | 1 | 6 |
| Total cell № | 28 | 117 | 134 | 104 | 44 | 103 | 68 | 118 | 103 | 137 | 86 | 363 |
| Average LDL (μm) | 234.2857 | 224.6154 | 205.0746 | 184.6154 | 222.7273 | 215.534 | 197.6471 | 179.661 | 223.301 | 216.9343 | 197.2093 | 178.4022 |

| Cell | UHZ-3 | | | | | | | |
|---|---|---|---|---|---|---|---|---|
| Experiments | 109-3 | | | | 112-3 | | | |
| LDL (μm) \ Speed | Slow | Normal | Fast 1 | Fast 2 | Slow | Normal | Fast 1 | Fast 2 |
| 360 | 0 | 0 | 0 | 0 | 0 | 0 | 1 | 0 |
| 320 | 5 | 3 | 6 | 19 | 5 | 12 | 7 | 23 |
| 280 | 7 | 5 | 0 | 21 | 10 | 7 | 16 | 10 |
| 240 | 23 | 28 | 19 | 105 | 23 | 32 | 23 | 128 |
| 200 | 15 | 20 | 45 | 53 | 18 | 35 | 49 | 72 |
| 160 | 9 | 25 | 7 | 176 | 9 | 31 | 7 | 171 |
| 120 | 2 | 5 | 2 | 38 | 2 | 5 | 3 | 34 |
| 80 | 0 | 0 | 2 | 14 | 0 | 0 | 3 | 21 |
| 40 | 0 | 1 | 4 | 15 | 0 | 1 | 6 | 17 |
| 0 | 0 | 0 | 2 | 10 | 0 | 0 | 3 | 5 |
| Total cell № | 61 | 87 | 87 | 451 | 67 | 123 | 118 | 481 |
| Average LDL (μm) | 225.5738 | 203.6782 | 197.2414 | 182.2616 | 226.8657 | 212.0325 | 206.4407 | 185.1975 |

**Supplemental Figure S2**

**Number and displaced length of cells separated by a microfluidic device. (a)** Magnified image of the exit of separation area, the position of inlet nozzle, and portions of lateral displacement. **(b)** Cell number and lateral displacement length (LDL) at each flow speed were shown. Separated cells were counted and the average LDL calculated (Materials and Methods). Three independent experiments were conducted in both U937 and UHZ-1, and two independent experiments in UHZ-3.

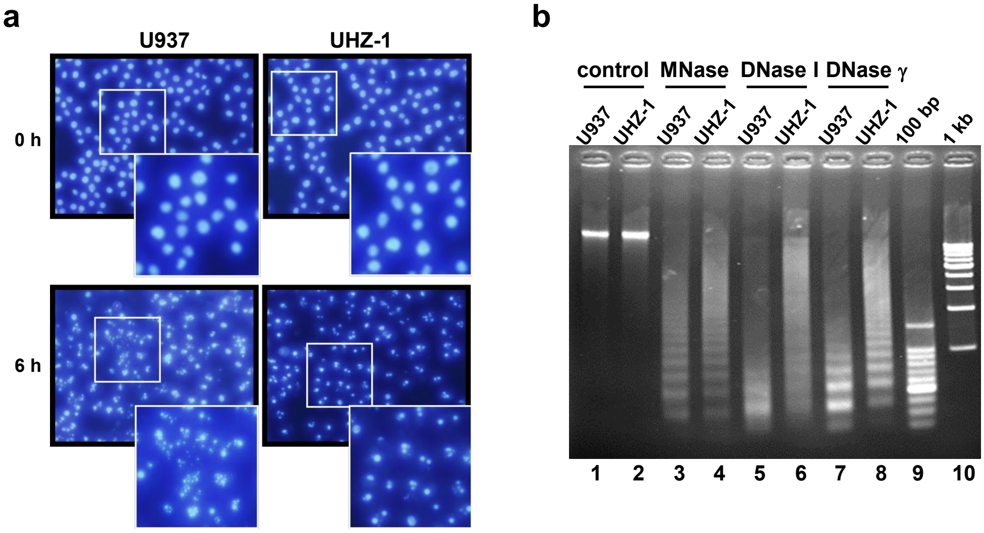


**Supplemental Figure S3**

**Increased histone H1 augments chromatin condensation.**

(**a**) More apoptotic chromatin condensation was detected in UHZ-1 than in U937. Apoptosis was induced by UV-irradiation and examined under a microscope at 6 h after the stimulation. Chromatin was visualized with Hoechst 33342. As the control, images at 0 h were shown. Magnified images were shown in insets. (**b**) The nuclei of UHZ-1 cells had more compact chromatin than those of U937 cells. The nuclei from U937 and UHZ-1 cells were incubated with DNases – MNase (lanes 3 and 4), DNase I (lanes 5 and 6), or DNase γ (lanes 7 and 8) – or without DNase (lanes 1 and 2). The purified DNA samples after incubation were analyzed by agarose gel electrophoresis. 100 bp: 100-bp DNA ladder marker. 1 kb: 1-kbp DNA ladder marker.
